# HDAC3 inhibitor RGFP966 enhances memory persistence in a biphasic manner and modulates NF-kB nuclear localization

**DOI:** 10.64898/2026.07.16.738883

**Authors:** Agustina Robles, Santiago D’hers, Mariana Feld, Arturo Romano

## Abstract

Nearly five decades ago, it was first observed that long-term memory consolidation requires waves of transcriptional activity and protein synthesis, with the first two waves occurring within hours after learning. While numerous studies have examined the effects of protein synthesis inhibitors, **how specific epigenetic regulators contribute to these waves remains poorly understood**. **Here, we aimed to determine whether HDAC3, a key modulator of memory, is involved in memory consolidation at these two time points through its potential relationship with the transcription factor NF-kB**, one of its deacetylation targets and a critical player in memory formation. Pharmacological inhibition of HDAC3 with RGFP966, either immediately or 6 hours after training, enhanced memory persistence in the NOR task in mice. Conversely, inhibition of NF-kB with BAY 11-7082 impaired memory at the same time points. Moreover, RGFP966 injection increased the nuclear proportion of NF-kB in the CA1 region of the hippocampus, suggesting that **HDAC3 inhibition is associated with changes in NF-kB nuclear localization**. To our knowledge, this study provides the first *in vivo* evidence that HDAC3 inhibition is associated with altered NF-kB localization during memory consolidation, extending previous observations from cell culture and electrophysiological studies in brain slices. **Although these findings support a functional association between HDAC3 inhibition and NF-kB signaling during consolidation, further experiments will be required to determine whether this relationship is mechanistically causal.**

## Introduction

Memory consolidation is the process by which newly acquired information is stabilized and stored for later retrieval. This process depends on gene expression and protein synthesis [1]. Epigenetic mechanisms play a central role in regulating gene expression during consolidation [2]. Among these, histone acetylation has been widely studied in the context of memory formation, as it increases DNA accessibility, thus promoting transcription [3]. Early studies found that inhibition of histone deacetylases (HDACs) improves LTP and contextual fear memory in rats [4], suggesting that increased histone acetylation promotes consolidation. Many studies later showed that HDAC inhibition promotes memory consolidation in different learning tasks in rodents, such as contextual fear conditioning (FC) [5, 6], inhibitory avoidance (IA) [7], Morris water maze (MWM) [8], Novel object recognition (NOR), and novel object location (NOL) [9–11]. On the contrary, experiments where acetylation was negatively affected by genetic manipulations, including mutations and deletions of the histone acetyltransferase (HAT) CBP, showed memory impairment [12, 13]. Interestingly, Stefanko and collaborators showed that inhibition of class I HDACs — the most abundantly expressed group of HDACs in the central nervous system [14, 15]— using Sodium Butyrate (NaBut) not only improves long-term memory at a 24-hour evaluation in the NOR task, but it also extends memory duration for at least one week [9]. Conversely, pharmacological HAT inhibition in the hippocampus does not affect NOR memory at 24 h but does impair memory persistence [16]. These findings suggest a role of epigenetic mechanisms in persistent forms of memory.

Among class I HDACs, HDAC3 has been proposed as a key negative modulator of memory [17]. It has been widely proven that abolishing its activity, either genetically or pharmacologically, improves memory formation [18–21], and its overexpression produces memory deficits [22, 23]. The mechanisms through which it downregulates memory include histone deacetylation in promoter regions of memory-related genes, such as *Bdnf*, *Fos*, *Nr4a2,* and *Npas4* [18–20, 24].

In the early 2000s, HDAC3 activity was linked to the NF-kB transcription factor induction and response duration in cell culture [25], **a relationship with potential relevance to memory, given NF-kB’s established role in memory consolidation and reconsolidation [26]**. Chen & Greene proposed that, upon entering the nucleus, the NF-kB p65 subunit is acetylated by CBP/p300, increasing its DNA binding and transcriptional activity. Conversely, HDAC3 removes the acetylation, promoting inhibitor protein IkB binding to NF-kB and subsequent nuclear expulsion, thus ending the signal [27]. There is evidence showing that NF-kB acetylation increases in the rat amygdala after an IA training [28], suggesting that this post-translational modification is relevant in the context of learning. Furthermore, it has been shown that HDAC3 inhibition restores synaptic tagging and capture in aged rats, and it does so through NF-kB activation [29]. These findings led us to hypothesize that HDAC3 negative modulatory activity might not be limited to decreasing histone acetylation levels, but could also be related to decreasing NF-kB presence in the nucleus.

One interesting feature of memory consolidation is that it involves recurrent waves of transcriptional activity. It was shown that there are two phases of protein synthesis in the rat hippocampus after an avoidance learning task, one around the time of training and the second around 6 hours later [30], and either transcription or translation inhibition at those time points produced memory impairment [31]. Similar results were obtained in other tasks, such as social recognition memory [32] and social fear memory [33] in mice. This biphasic nature of the molecular processes involved in memory formation is also present in the crab *Neohelice granulata*, where NaBut injection immediately or 6 hours after training improved memory retention [34]. Additionally, two peaks of PKA [35] and NF-kB activity [36] were found at the same time points mentioned above. Considering that NF-kB biphasic activity has already been demonstrated in invertebrates and given the key role of HDAC3 in regulating memory-related gene expression, we sought to determine whether inhibiting their activity affects memory in mice following a similar temporal pattern.

This study aimed to investigate the temporal pattern of HDAC3 and NF-kB involvement during the consolidation of a persistent memory in mice using the NOR task. To that end, we performed pharmacological experiments where either HDAC3 or NF-kB was inhibited at different times after training and then evaluated memory retention one week later. We also performed molecular and immunostaining experiments to further evaluate the effect of HDAC3 inhibition on the NF-kB pathway during memory formation.

## Materials and Methods

### Animals

Male CF1 (FVET - UBA) mice of 8 weeks of age were used for the behavioral experiments involving systemic administration of the drug RGFP966 and memory evaluation. The remaining experiments were conducted on 8-week-old male and female C57BL/6 mice (IMEX - ANM - CONICET). Animals were housed in the Institute of Physiology, Molecular Biology and Neuroscience (IFIBYNE) animal facility under controlled conditions: 2-4 animals per cage, 24°C, 12/12 h light/dark cycle with lights on at 7 a.m. Food and water were provided *ad libitum*. Every procedure was in accordance with the guidelines approved by the Institutional Commission for the Care and Use of Laboratory Animals (CICUAL – FCEN – UBA), protocol n°153.

### Behavioral tasks

#### Novel Object Recognition

The experimental arena consisted of a white box (31 x 23 x 31 cm; l x w x h) with the floor covered by sawdust. Before the start of the experiment, animals were handled for 2-3 days. The first two sessions consisted of habituation to the empty arena and took place on consecutive days, lasting 5 and 15 min, respectively. Twenty-four hours later, on training day, animals were exposed to two identical objects. Weak training sessions lasted 3 min for C57BL/6 mice [16] and 1 min for CF1. Strong training sessions lasted 15 min. On testing day, animals were faced with one familiar object and a novel object for 5 min. Objects were cleaned thoroughly between animals with 70% ethanol, and sawdust was rearranged to avoid any scent trails. The objects were a 50 ml glass beaker (turned upside down) and a LEGO block of similar size, which were fixed to the bottom of the arena with Velcro. Every session was recorded with a Genius Ecam 800 Full HD webcam for later analysis. Exploration time was scored manually using the software AnyMaze and was defined as the times when the animal was sniffing or touching the objects, or had its head oriented towards the object at a maximum distance of 1 cm from its perimeter. Discrimination index (DI) was calculated as the ratio of time spent exploring the novel object to total exploration time and expressed as a percentage.

#### Fear Conditioning

The arena consisted of an acrylic box (24.5 x 24.5 x 42 cm; l x w x h) with a metallic grid on the floor for shock delivery, placed inside a white wooden box equipped with a speaker and dimmable light. Animals were handled for 3 consecutive days before the start of the experiment. On training day, animals were placed inside the arena, and after 2 min of habituation, they received either 1 or 3 trials consisting of a tone (10 sec, 80 dB) co-terminating with a foot-shock (1 sec, 0.6 mA). Animals were tested for context memory in the training context (context A) in the absence of either tone or shock, for 5 min. The next day, they were tested with the tone for cued memory in a different context (context B). For context B, we used the same acrylic box, with the metallic grid covered with a white plastic floor, and a green acetate sheet placed inside the arena in a semi-circular arrangement to change the color and shape of the walls. A small piece of filter paper was impregnated with vanilla extract scent and placed inside the wooden box to change olfactory cues, and the lights were dimmed. Contexts were thoroughly cleaned between animals with either 70% ethanol (for context A) or 2% acetic acid (for context B). Freezing was scored manually every 5 seconds, and the percentage was calculated by dividing freezing events by the total number of events.

### Surgery

Animals were anesthetized with Isoflurane (3% for induction, 1.5-2% for maintenance). Stereotaxic surgery was then performed, and cannulae were implanted in the dorsal hippocampus as previously described [37]. Briefly, after placing the animals in the stereotaxic frame, the skull was exposed, and the bregma was identified using small amounts of hydrogen peroxide and cotton swabs. Coordinates were determined relative to the bregma (A-P: -1.9 mm; M-L: ±1.2 mm). After drilling the holes, cannulae were implanted to a depth of 1.2 mm and secured to the skull with dental acrylic.

To minimize pain, animals received a subcutaneous injection of meloxicam (5 mg/kg) immediately after surgery, and tramadol (100 mg/ml) was provided in the drinking water (1 drop/100ml) for the next 48 h. Behavioral procedures started 7 days after surgery to allow proper recovery.

### Drugs

For experiments involving systemic administration, animals received i.p. injections of class I HDAC inhibitor NaBut (Sigma; 1.2 g/kg) [9], HDAC6 specific inhibitor Tubastatin A (TubA; Cayman Chemical, Ann Arbor, MI; 40 mg/kg) [38], or HDAC3 specific inhibitor RGFP966 (Sigma; 10 mg/kg) [18] dissolved in vehicle, composed of 15% DMSO, 10% polyethylene glycol 400 and 2% Tween-20 in sterile saline solution.

For intra-hippocampal infusions, NF-kB inhibitor BAY 11-7082 was dissolved in DMSO to a 65 mM concentration (adapted from Zhang et al., 2018) [39]. Animals received bilateral injections of 0.5 μl on each side through a device consisting of a 5 μl Hamilton syringe connected to a 30 G needle. Infusions lasted for 30 s, followed by a 30 s pause before needle withdrawal to prevent reflux.

### Protein extracts and Western blot assay

Animals were euthanized by cervical dislocation, and the hippocampus was removed. Tissue was homogenized in 250 μl of Buffer A (10 mM HEPES pH 7.9, 10 mM KCl, 1.5 mM MgCl_2_,1 mM DTT, 1 μg/ml pepstatin A, 10 μg/ml leupeptin, 0.5 mM PMSF, 10 μg/ml aprotinin, 5 mM sodium butyrate, 1 mM sodium orthovanadate, and 50 mM NaF) with 15 strokes using a glass-glass Dounce homogenizer with a type B pestle. Homogenates were centrifuged for 15 min at 1000 g and 4°C. Supernatant was collected and stored at -20°C until use. Protein concentration was measured using a BCA Protein Assay (Thermo Scientific) following the manufacturer’s instructions.

For the western blots, samples were mixed with loading buffer, heated at 95°C for 5 min, and immediately placed on ice. Twenty μg of total protein was loaded in a 12.5% denaturing acrylamide gel for electrophoresis. Proteins were then electrotransferred to a polyvinylidene fluoride (PVDF) membrane. After blocking, membranes were incubated with primary antibodies overnight at 4°C and then with secondary antibodies at room temperature for 1 h. Immunoblotting was performed with the following antibodies: [Z]-p65 (1:1000, Cell Signaling – 6956), [Z]-p65ac K310 (1:1000, Abcam – ab218533), [Z]-Erg1 (1:1000, Santa Cruz – sc-110), [Z]-GAPDH (1:1000, Santa Cruz – sc-32233), [Z]-Mouse HRP (1:10000, Cell Signaling – 7076), [Z]-Rabbit HRP (1:5000, Santa Cruz – sc2357). Detection was performed using Luminol reagent (0.1 M Tris pH 8.8, 1.25 mM Luminol, 0.198 mM p-coumaric acid, 0.0096% hydrogen peroxide), and images were acquired with an Amersham Imager 600.

### Brain slices and immunofluorescence

Animals were deeply anesthetized (ketamine 150 mg/kg; xylazine 10 mg/kg) and then transcardially perfused with 20 ml saline solution followed by 30 ml of paraformaldehyde (PFA) 4% in PB 0.1 M. Brains were extracted, post-fixed in PFA 4% for 24 h, and then transferred to a 30% sucrose solution in PB 0.1 M for at least another 24 h. Intact brains were maintained at 4°C. Coronal sections of 40 μm were obtained using a Leica Microm HM 505 N cryostat and collected in cryopreserving solution (3 vol. glycerol, 3 vol. ethylene glycol, 4 vol. PB 0.2 M) stored at -20°C until use.

Slices corresponding to the dorsal hippocampus and the perirhinal cortex were selected (between -1.58 and -2.06 mm according to Paxinos & Franklin) [40]. After washing with PBS, slices were incubated in sodium citrate 10 mM, 0.05% Tween-20, pH = 6, for 25 min at 85°C for antigenic retrieval. Then, they were washed again, followed by blocking with 2.5% BSA in PBS for 15 min at 37°C. Incubation with primary antibody [Z]-p65 (1:200, Cell Signaling – 6956 or 1:100, Cell Signaling - 8242) diluted in PBS, 0.05% Triton, and 0.5% BSA occurred overnight at 4°C. Secondary antibody ([Z]-Mouse Alexa 488, 1:1000, Invitrogen – A-11001; [Z]-Rabbit Alexa 647, 1:1000, Invitrogen – A-21245) was diluted in PBS with 0.1% Triton and 0.5% BSA, and incubation lasted 90 min at room temperature. Nuclei were stained with DAPI (1:1000, HelloBio cat. HB8199) dissolved in PBS. Finally, slices were mounted with Mowiol and kept protected from light. Images were acquired with a Zeiss LSM900 confocal microscope using an objective lens 40x NA 1.3. Stacks contained 10 – 20 images with a voxel depth of 0.5 μm.

### Immunofluorescence Image Analysis

To quantify fluorescence, we first performed a 3D segmentation of the nuclei using Cellpose (v3.0) [41] by filtering and training a customized model starting from one of the pre-trained models [42, 43]. Once the 3D masks of the nuclei were obtained, the fluorescent signal was quantified using CellProfiler (v4.2.8) [44]. To measure the size and number of the particles inside the nucleus, another mask was created, isolating these objects and associating them with their corresponding nucleus (also using CellProfiler). The model and pipelines used can be found at (https://github.com/AgusRobles/Confocal-image-analysis). Representative images show one slice of the z-stack.

The number of p65-immunopositive cells was counted manually using Fiji/ImageJ (v2.16.0) [45]. Representative images were generated using a z-projection of the sum of intensities of the z-stack.

We included 4 brain slices from each animal in every experiment.

### Statistical Analysis

Plots and statistical analyses were performed using R (v4.4.1) [46] and RStudio (v2026.1.1.403) [47]. Data were analyzed using linear mixed models (LMMs) followed by ANOVA or generalized mixed models (GLMMs) followed by Deviance analysis (type II Wald chi-square tests). *A priori* planned comparisons between treatments within each level of the remaining factors were performed using the Tukey method with the “emmeans” package [48]. Proportions (DI and fluorescence ratio) were modelled using a Beta distribution and a logit link, freezing percentage with a binomial distribution, and speckle counts per nucleus with a quasi-Poisson distribution. GLMMs were fitted with the “glmmTMB” package (v1.1.14) [49, 50]; model assumptions were checked using “DHARMa” (v0.4.7) [51]; and overdispersion for the quasi-Poisson model was additionally verified using the “performance” package [52]. Experiments with repeated observations from the same animal included the animal ID as a random effect to account for the non-independence of the measurements. **For immunofluorescence experiments, brain sections were modeled as repeated observations nested within animals to avoid pseudoreplication.** Experiments including animals from both sexes were analyzed with sex as a random effect. Intraclass correlation coefficients (ICC) were calculated for mixed-effects models using the “performance” package [52] to quantify the proportion of variance attributable to the random effects.

## Results

### HDAC3 inhibition promotes memory persistence in a biphasic manner in the NOR task, but does not affect fear memory

Experiments using the general HDAC class I inhibitor NaBut in mice show that systemic administration promotes memory persistence for at least one week after training in the NOR task [9]. In parallel, accumulating evidence identifies HDAC3 as a negative memory modulator, and its inhibition has proved to promote consolidation of different types of memory, such as NOL, NOR, and both cued and contextual fear conditioning [17–19, 53]. Based on these findings, we asked whether systemic HDAC3 inhibition could promote memory persistence in the NOR task in mice. We performed an experiment where four groups of male mice received a weak training (wTR) in the NOR task and, immediately after, were injected with the HDAC3-specific inhibitor RGFP966, NaBut, TubA (a specific HDAC6 inhibitor), or vehicle. Given that NaBut has already been shown to enhance memory persistence, we used it as a positive control. Considering that both NaBut and RGFP966 target nuclear HDACs, and that HDAC6 -a cytosolic HDAC- has been implicated in memory processes [38, 54], we included TubA to assess the effect of systemic HDAC6 inhibition on memory persistence. Memory was assessed 1 week after training (Fig. 1a) and found an overall significant effect of the treatment on the DI during the testing session (χ^2^_(3)_= 22.348, p < .001). Both NaBut- and RGFP966-injected animals showed a significant increase compared to vehicle-injected controls (Fig. 1b; DI_VEH_ = 52.9 ± 3.28%; DI_NaBut_ = 64.8 ± 2.75%; DI_RG_ = 68.4 ± 1.39%; NaBut vs Veh: p < .05; RGFP966 vs veh: p < .001), indicating better memory performance. On the other hand, animals treated with TubA showed no differences compared to controls (Fig. 1b; DI_TubA_ = 54.5 ± 3.68%; TubA vs Veh: p > .05). To assess recognition memory for the familiar object, we analyzed the time spent exploring each object and performed a two-way ANOVA with object familiarity (familiar or novel) and treatment (vehicle or drug) as factors. We found a significant interaction between treatment and familiarity (F_(3, 33)_ = 10.265, p < .001), and *post-hoc* comparisons showed that both NaBut- and RGFP966-injected animals explored significantly more the novel than the familiar object (Fig. S1a; p < .001), while TubA and vehicle groups explored both objects equally (Fig. S1a; p > .05). Although systemic RGFP966 administration has been previously shown to facilitate memory consolidation in the NOR task when tested at 24 h [18], our results indicate that memory enhancement persists for at least one week. Our findings also confirm the effect previously reported for NaBut and show that HDAC6-specific inhibition does not promote memory persistence in the NOR task.

**Fig. 1.**
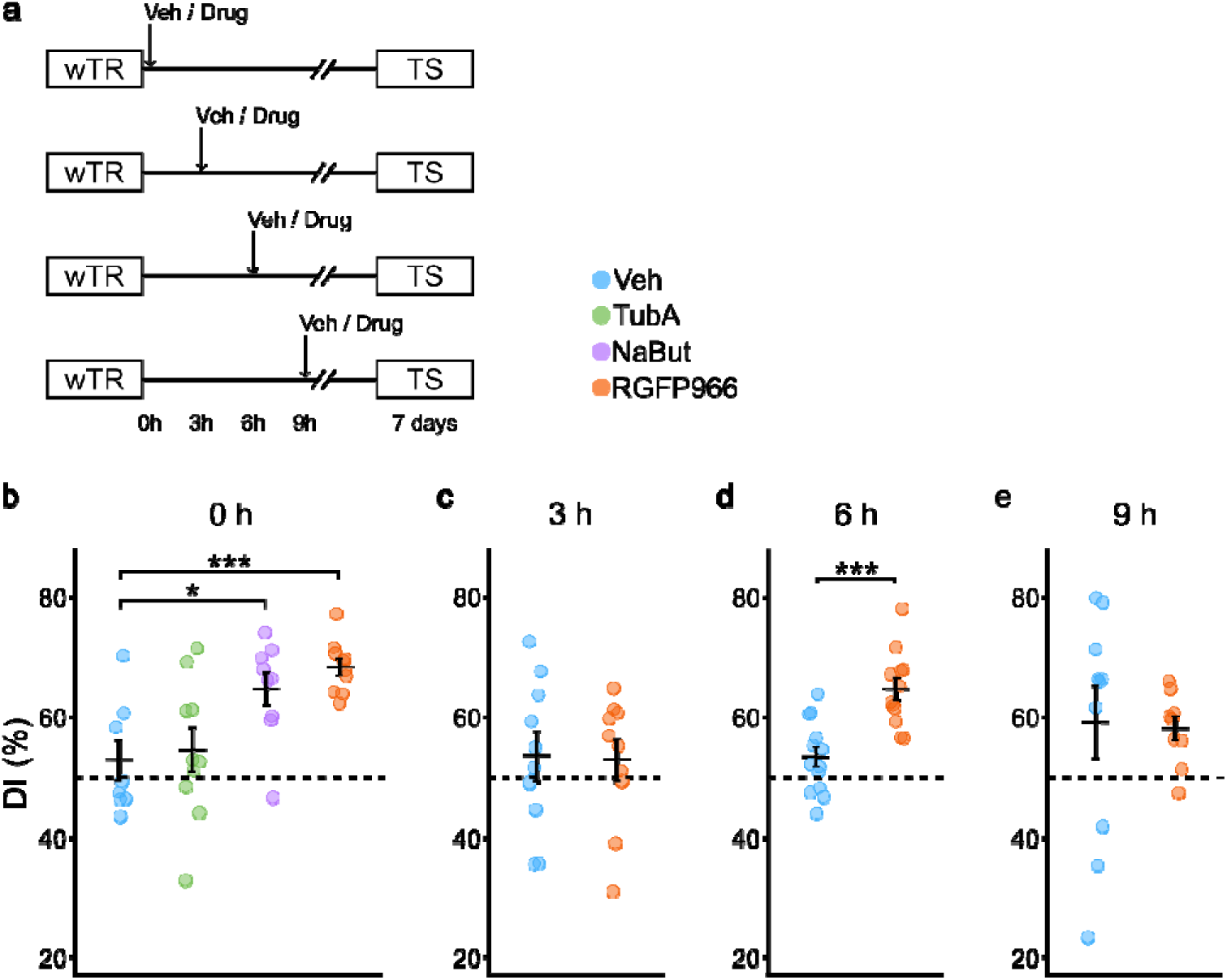
HDAC3 inhibition with RGFP966 immediately or 6 h after training enhances NOR memory persistence. General HDAC inhibition with NaBut immediately after training also enhances memory persistence, while HDAC6 inhibition (TubA) does not. (**a**) Experimental design. Animals received a weak training session and were then injected i.p. with either drug or vehicle at different time points: (**b**) immediately (n_VEH_ = 8, n_TubA_ = 10, n_NaBut_ = 9, n_RGFP966_ = 10; contrasts against VEH corrected by Dunnet’s method), (**c**) 3 h (n_VEH_ = 10, n_RGFP966_ = 10), (**d**) 6 h (n_VEH_ = 13, n_RGFP966_ = 12), or (**e**) 9 h (n_VEH_ = 10, n_RGFP966_ = 9) after training. Memory was assessed 1 week later. Mean DI (%) ± SEM are shown in black. Individual values are colored by treatment. * p < .05, *** p < .001; GLM, type II Wald’s test

Considering the evidence stating that there are two waves of gene expression and protein synthesis shortly after learning during consolidation, we asked whether HDAC3 inhibition at other time points after training could also affect memory persistence. To further investigate the effect of the drug during the consolidation window, we carried out three independent experiments with drug or vehicle injections at either 3, 6, or 9 h after training (Fig. 1a). We found that drug injection 6 h after training also improved memory as evidenced by the significant increase in the DI (Fig. 1d; DI_VEH_ = 53.4 ± 1.66%; DI_RG_ = 64.8 ± 1.83%; χ^2^_(1)_ = 22.419, p < .001). Exploration times confirmed this result, as only the animals in the RGFP966 group spent significantly more time exploring the novel object (Fig. S1c; treatment-by-familiarity: F_(1, 23)_ = 15.6549, p < .001; RGFP966: familiar – novel p < .001; VEH: familiar – novel p > .05). On the other hand, injections 3 or 9 h after training did not alter DIs whatsoever, as no significant differences were observed between drug- and vehicle-injected animals (Fig. 1c; 3 hs: DI_VEH_ = 53.6 ± 4.01%; DI_RG_ = 53 ± 3.38%; χ^2^_(1)_ = 0.0256, p > .05; Fig. 1e; 9 hs: DI_VEH_ = 59.2 ± 6.05%; DI_RG_ = 58.1 ± 1.99%; χ^2^_(1)_= 0.0656, p > .05). Regarding exploration times, in the 3 h experiment we found no interaction (F_(1, 18)_ = 0.0353, p > .05), and no significant main effects of either treatment (F_(1, 18)_ = 0.1364, p > .05) or familiarity (F_(1, 18)_ = 1.0775, p > .05), indicating that there is no memory of the familiar object (Fig. S1b). Curiously, although DIs did not differ, animals injected 9 h after training showed higher exploration of the novel object relative to the familiar one, irrespective of the treatment (treatment-by-familiarity: F_(1, 17)_ = 0.185, p > .05; treatment: F_(1, 17)_ = 0.1026, p > .05; familiarity: F_(1, 17)_ = 8.9323, p < .01). Comparisons between time spent exploring the familiar vs the novel object showed a significant difference (Fig. S1d; p < .01) indicating that there is memory of the familiar object that was not altered by the drug. These results show that HDAC3 inhibition by systemic administration of RGFP966 can promote persistence of a weak memory, and that this effect has a biphasic window of action.

Given the marked effect of RGFP966 found in the NOR task, and previous work showing that catalytically dead HDAC3 expression in the dorsal hippocampus or amygdala subnuclei promotes contextual and auditory fear memory consolidation [19], we asked whether pharmacological HDAC3 inhibition could produce the same effect. We used a Fear Conditioning task with three groups of male and female mice: two received a weak training session (wTR, 1 pairing tone-shock), immediately followed by an i.p. injection of either drug or vehicle, while the third group received a strong training (sTR, 3 pairings tone-shock) followed by a vehicle injection as a positive control for learning. Testing sessions occurred at 24 h, 1, and 2 weeks after training, with context and tone memory assessed on consecutive days (Fig. S2a). For contextual fear, we found a significant group-by-session interaction (χ^2^_(4)_= 14.258, p < .01), with the sTR group differing significantly from both wTR groups, while the wTR groups did not differ from each other indicating no drug effect (Table S1; Table S2) Contextual freezing decreased over time in all groups, more steeply in the sTR group, likely reflecting a floor effect in the wTR groups (Fig. S2b-d).

For cued fear, we found significant main effects of group (χ^2^_(2)_= 8.2678, p < .05) and testing session (χ^2^_(2)_= 76.2629, p < .001), but no interaction (X^2^_(4)_= 1.4368, p > .05). *Post-hoc* comparisons showed significant differences between the strong and weak-trained groups only at the 8 d test (Fig. S2f), with no effect of the drug at any session (Fig. S2e-g; Table S1; Table S2).

Minute-by-minute analysis of freezing within sessions gave the same pattern as total freezing, ruling out the possibility that the drug could be affecting the dynamics of the freezing behavior during testing (Fig. S2). Total distance travelled and mean speed did not differ between groups (data not shown), arguing against an effect on non-freezing fear behaviors.

Overall, we found no effect of systemic HDAC3 inhibition on either contextual or cued fear memory consolidation. **However, results from the later testing sessions should be interpreted with caution, as previous retrieval may have modified the memory through reconsolidation.**

### Systemic HDAC3 inhibition decreases Zif268 levels in the hippocampus

Experiments in cell culture have shown a functional link between HDAC3 and NF-kB. Upon entering the nucleus, NF-kB is acetylated by CBP/p300, preventing binding of its inhibitor IkB, and sustaining activation. Conversely, HDAC3 has been proposed to reverse this, promoting IkB binding and subsequent cytosolic export [25]. Considering that NF-kB activity has been directly linked to memory consolidation [26], we hypothesized that HDAC3-mediated cytoplasmic translocation of NF-kB could be a mechanism underlying the behavioral enhancement observed. To test this, we used the same protocol that produced memory enhancement (Fig. 2a). Consequently, two groups of mice received a wTR in the NOR task, followed immediately by i.p. injection of either RGFP966 or vehicle. Instead of evaluating memory retention, mice were sacrificed 45 min later, the reported peak of NF-kB activity after training [36, 55], to obtain hippocampal protein-enriched extracts that were analyzed by western blot (Fig. 2b). We found no differences in p65 K310 acetylation or total p65 expression levels (Fig. 2c; p65ac_VEH_ = 1 ± 0.24; p65ac_RG_ = 0.73 ± 0.15; F_(1, 13)_ = 1.4802, p > .05; Fig. 2d; p65_VEH_ = 1 ± 0.11; p65_RG_ = 0.97 ± 0.13; F_(1, 13)_ = 0.0397, p > .05). Interestingly, p65 acetylation showed an ICC of 0.533 indicating that 53.3% of the variability is explained by the sex, pointing to a potential sexual dimorphism, with vehicle-injected males displaying lower acetylation levels than females (Fig. 2c). We also measured Zif268 protein level, an immediate early gene regulated by NF-kB and HDAC3 [56]. Zif268 was significantly lower in the drug-treated group compared to vehicle-injected animals (Fig. 2e; Zif268_VEH_ = 1 ± 0.27; Zif268_RG_ = 0.48 ± 0.13; F_(1, 13)_ = 5.4564, p < .05). Furthermore, a high ICC (58.4%) also suggested a sexual dimorphism, with the observed decrease occurring only in males, while females showed similarly low Zif268 levels in both experimental conditions. Notably, a similar pattern of Zif268 expression was observed in naïve animals (Zif268_VEH_ = 1 ± 0.28 (n = 8), Zif268_RG_ = 0.22 ± 0.1 (n = 5), F_(1, 10)_ = 5.4673, p < .05, ICC = 42.8%), indicating a sex-dependent effect of the drug.

**Fig. 2.**
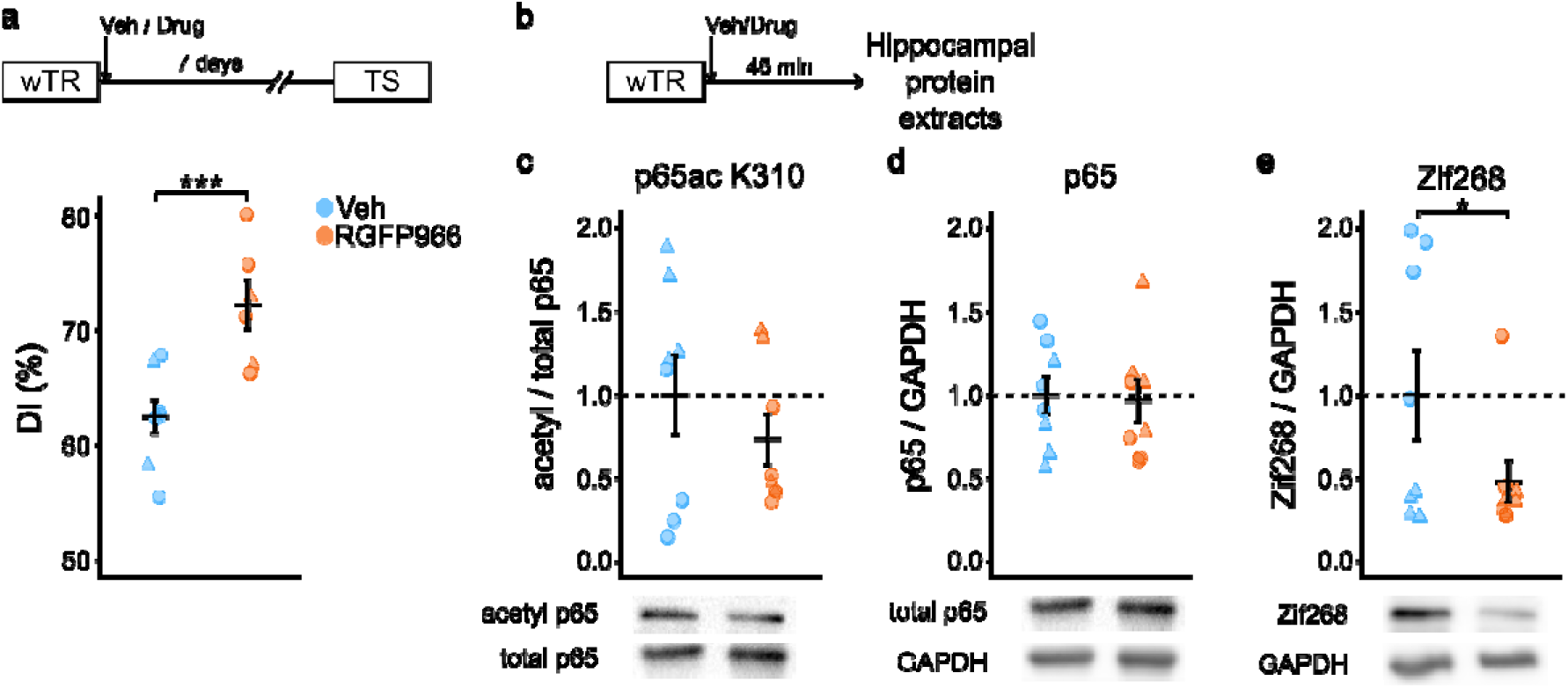
HDAC3 systemic inhibition with RGFP966 does not affect p65 acetylation levels, but decreases Zif268 levels in hippocampus extracts. (**a**) DI (%) of animals receiving a wTR followed by an i.p. injection of either drug or vehicle, and evaluated 1 week later (n_VEH_ = 8, n_RG_ = 6). (**b**) Experimental design. Animals received a wTR in the NOR task immediately followed by an i.p. injection of drug or vehicle. Hippocampal protein extracts were obtained 45 min after injection. (**c-e**) Quantification of band intensities of (**c**) acetylp65/total p65, (**d**) p65, and (**e**) Zif268 relative to GAPDH (n_VEH_ = 8, n_RGFP966_ = 8). Representative western blots are shown at the bottom. Mean ± SEM are shown in black. Individual data points are colored by treatment. ▴: females, □: males. * p < .05, *** p < .001; linear mixed-effects model

### Systemic RGFP966 administration after training increases the proportion of nuclear p65 in CA1

In the previous experiment, we studied p65 acetylation in response to HDAC3 inhibition **in whole-hippocampus extracts and found no effect of the drug. However, this approach cannot resolve differences between hippocampal subregions and did not include other relevant structures such as the perirhinal cortex (Prh). We therefore turned to immunofluorescence, which allowed simultaneous assessment of p65 nuclear localization across CA1, CA3, and dentate gyrus (DG) subregions of the dorsal hippocampus, as well as the Prh, a region crucial to object recognition** [57]. Given our hypothesis that HDAC3 activity promotes NF-kB nuclear export, we predicted that HDAC3 inhibition would favor NF-kB nuclear localization.

Two groups of mice received a wTR in the NOR task followed immediately by an i.p. injection of either RGFP966 or vehicle. Forty-five min later, animals were perfused, and coronal brain sections were immunostained for p65 (Fig 3a). We analyzed the ratio of nuclear-to-total fluorescence intensity across regions, finding no significant treatment-by-region interaction (χ^2^_(3)_= 1.3058, p > .05), but significant main effects of treatment (χ^2^_(1)_= 5.6711, p < .05) and region (χ^2^_(3)_= 181.5796, p < .001). *Post-hoc* contrasts showed a higher proportion of nuclear p65 in CA1 in RGFP966-treated animals compared to vehicle-treated controls (Fig. 3b (left), c; VEH = 0.16 ± 0.01; RG = 0.18 ± 0.01; p < .05), whereas no changes were observed in CA3, DG (Fig. S3b-e), or Prh (Fig. 3d (left), e). **These results are consistent with the possibility that HDAC3 inhibition promotes NF-kB nuclear localization as a candidate mechanism for the observed memory enhancement, though given the modest sample size this should be interpreted with caution.**

**Fig. 3.**
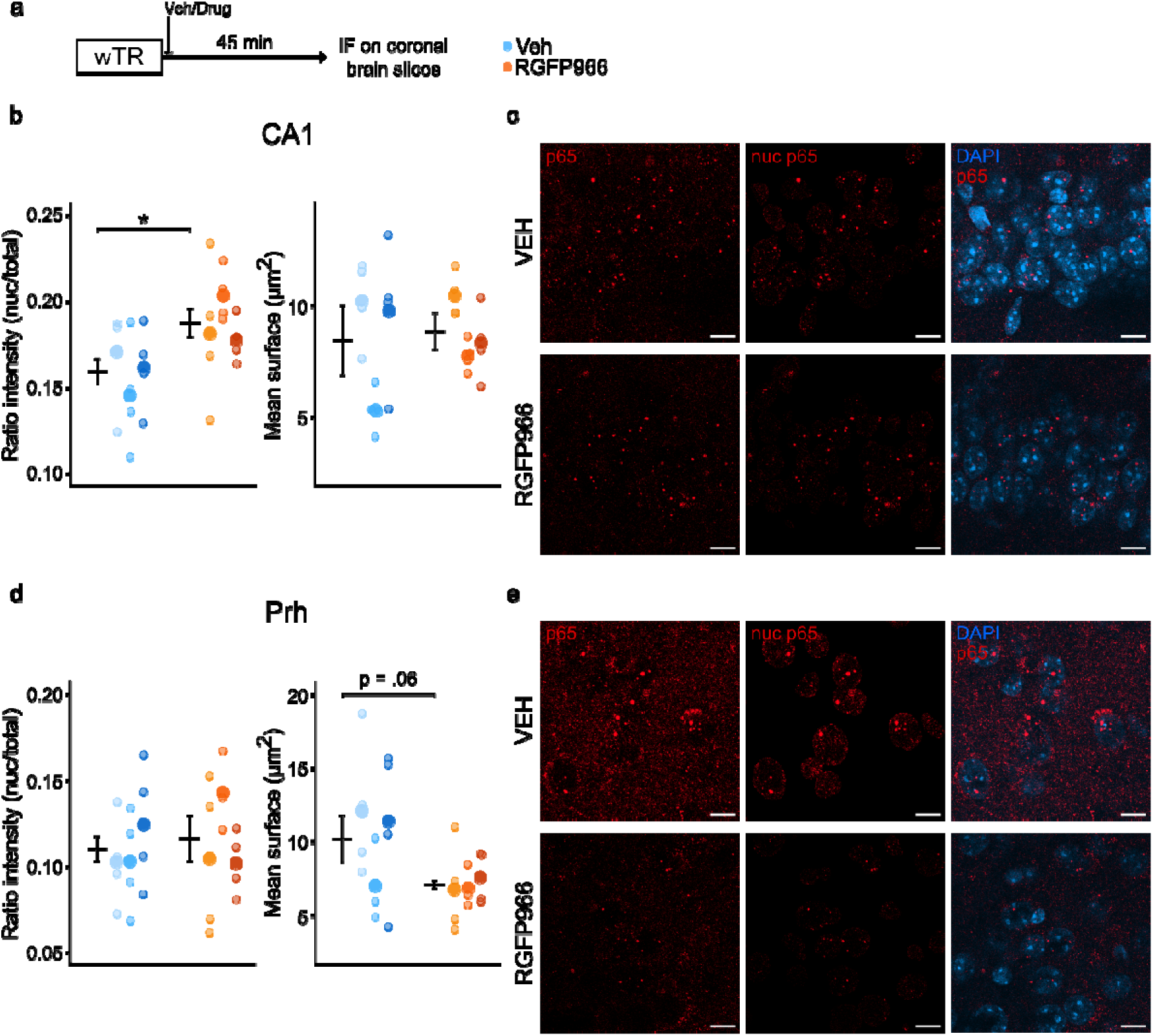
Systemic administration of RGFP966 after training increases the proportion of nuclear p65 in the CA1 region of the hippocampus and changes the signal distribution in the Perirhinal cortex (Prh). (**a**) Experimental design. Animals received a wTR in the NOR task, followed by i.p. injection of either drug or vehicle. Forty-five min later, p65 total fluorescence was measured in CA1, DG, CA3, and Prh brain regions. (**b, d**) (Left) Fluorescence intensity expressed as the proportion of nuclear signal (nuclear p65 / total p65; GLM, type II Wald’s test), and (right) 3D surface area of the p65 speckles inside the nucleus (two-way ANOVA) in (**b**) CA1 and (**d**) Prh. (**c, e**) Confocal images of (**c**) CA1 and (**e**) Prh. Mean ± SEM are shown in black. Individual data points are colored by treatment. Data from each animal are displayed in separate columns; small dots represent individual brain slices, and the large dot indicates the mean for that animal (n_VEH_ = 3, n_RGFP966_ = 3). Scale bar = 10μm. * p < .05; simple pairwise comparisons Tukey-adjusted

We also noted that p65 signal formed speckle-like nuclear aggregates and asked whether their number or size differed between treatments, which could suggest a possible redistribution of NF-kB inside the nucleus. Using a 3D mask to isolate speckles within each nucleus, we found a significant effect of treatment-by-region interaction in speckle number (χ^2^_(3)_= 118.349, p < .001). However, *post-hoc* comparisons showed no significant differences between RGFP966- and vehicle-treated groups in any individual region (Fig. S3f-i).

Next, we evaluated the size of the speckles, measuring both volume and surface area. Since the speckles were mostly spherical, both measures yielded very similar results; therefore, only surface area is shown. A two-way ANOVA showed no significant treatment-by-region interaction (F_(3, 64)_ = 2.4286, p = .07) or treatment main effect (F_(1, 4)_ = 0.7157, p > .05). Only a significant effect of the region was found (F_(3, 64)_ = 9.8023, p < .001). No subregion of the hippocampus showed treatment differences (Fig. 3b (right), Fig. S3b, d); however, there was a marked decrease in the size of the speckles in the Prh cortex of RGFP966-injected animals, although it was not statistically significant (Fig. 3d (right); VEH = 10.5 ± 1.41; RG = 7.15 ± 0.64; p = .06). Descriptive statistics and treatment comparisons for all variables are summarized in Table S3.

In conclusion, systemic HDAC3 inhibition was associated with an increase in the proportion of nuclear p65 in CA1 during object recognition memory consolidation, and may alter its distribution inside the nucleus in the Prh. These findings suggest a possible mechanism underlying the drug’s effect on memory persistence.

### NF-kB intra-hippocampal inhibition negatively affects memory persistence in a biphasic manner and decreases p65^+^ cells in CA1

Previous research from our lab showed that NF-kB inhibition with Sulfasalazine or kB-decoy (a competitive binding probe designed to sequester the transcription factor) during consolidation and reconsolidation impairs memory across different tasks in mice [16, 37]. Considering the two critical time windows identified for RGFP966 and its effect on p65 distribution in the brain, we investigated the time course of NF-kB pathway inhibition during object recognition memory consolidation, using BAY 11-7082, a structurally distinct inhibitor whose acute effects on memory in mice have not, to our knowledge, been previously tested.

As it was unclear whether BAY 11-7082 crosses the blood-brain barrier, we performed a preliminary experiment with systemic administration: two groups of animals received a strong 15-minute training (sTR) session followed by i.p. Injection of BAY or vehicle, and memory was evaluated 1 week later. No significant differences in the DI were observed between groups, with both groups showing higher levels of novel object exploration compared to the familiar object (Fig. S4). These results suggest that systemic BAY administration does not affect memory under our experimental conditions and prompted us to assess localized hippocampal administration.

Mice were implanted with bilateral cannulae in the dorsal hippocampus and, one week after surgery, both groups received a sTR in the NOR task, immediately followed by a localized intra-hippocampal injection of either BAY or vehicle (Fig. 4a). When tested one week later, BAY-injected animals displayed significantly lower Dis compared to vehicle-injected controls (Fig. 4b; DI_VEH_ = 73.4 ± 3.17%; DI_BAY_ = 65.1 ± 2.4%; χ^2^_(1)_ = 5.4637, p < .05). Exploration time analysis showed no significant treatment-by-familiarity interaction (χ^2^_(1)_ = 0.7365, p > .05), but significant main effects of treatment (χ^2^_(1)_ = 9.8999, p < .01) and familiarity (χ^2^_(1)_ = 46.3924, p < .001), with both groups exploring the novel object more (Fig. S5a; p < .001). Thus, while all animals remembered the familiar object, hippocampal NF-kB inhibition immediately after training altered memory performance, producing a decrease in the DI.

**Fig. 4.**
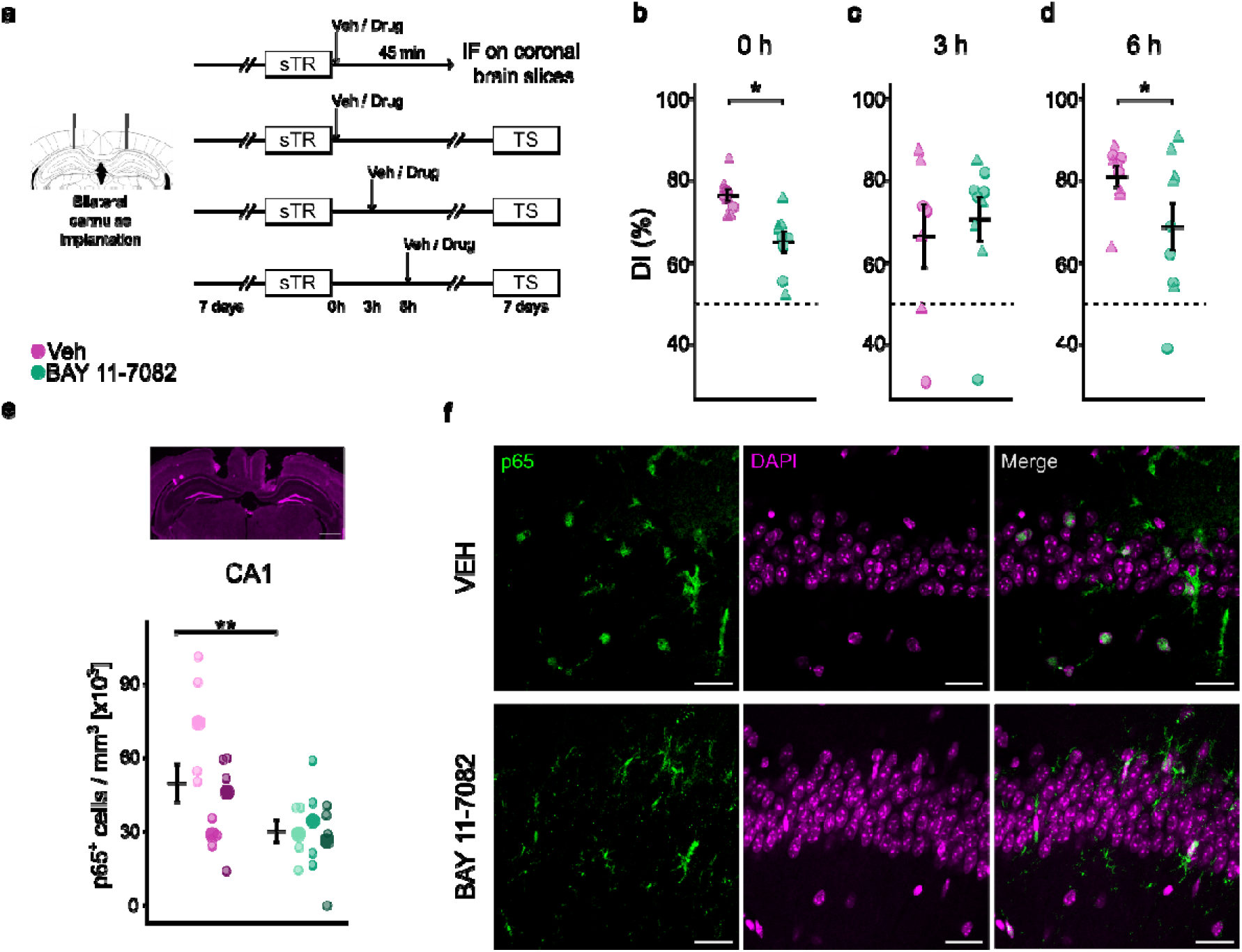
Intra-hippocampal NF-kB inhibition with BAY 11-7082 immediately or 6 h after training negatively affects memory persistence in the NOR task and decreases p65^+^ cells in CA1. (**a**) Experimental design. Animals underwent cannulae implantation surgery and, 7 days later, received a sTR in the NOR task followed by intra-hippocampal injections of drug or vehicle at different times. (**b-d**) DI (%) of animals receiving intra-hippocampal injections (**b**) immediately (n_VEH_ = 10, n_BAY_ = 9), (**c**) 3 h (n_VEH_ = 8, n_BAY_ = 10), or (**d**) 6 h (n_VEH_ = 9, n_BAY_ = 9) after training (GLMM, type II Wald’s test). (e) Quantification of the number of positively labeled cells per mm^3^ in CA1 (two-way ANOVA, simple pairwise comparisons, Tukey-adjusted). (**f**) Confocal images of brain slices stained against p65 (n_VEH_ = 3, n_BAY_ = 3). Scale bar = 25μm. Mean ± SEM are shown in black. Individual data points are colored by treatment. Data from each animal are displayed in separate columns; small dots represent individual brain slices, and the large dot indicates the mean for that animal. ▴: females, □: males. * p < .05, ** < .01

We next tested BAY administration at 3 and 6 h after training, to determine whether NF-kB inhibition shows the same biphasic time window seen with RGFP966 (Fig. 4a). At 3 h, BAY had no effect on the DI (Fig. 4c; DI_VEH_ = 66.3 ± 7.6%; DI_BAY_ = 70.6 ± 5.34%; χ^2^_(1)_= 0.1464, p > .05), and both groups explored the novel object significantly more than the familiar one (Fig. S5b; treatment-by-familiarity: F_(1, 14)_ = 0.3725, p > .05; treatment: F_(1, 14)_ = 0.042, p > .05; familiarity: F_(1, 14)_ = 19.9022, p < .001; familiar vs novel: VEH p < .05, BAY p < .01). Interestingly, when animals were injected 6 h after training, the BAY-injected group showed significantly lower DI than controls (Fig. 4d; DI_VEH_ = 80.9 ± 2.54%; DI_BAY_ = 68.7 ± 5.82%; χ^2^_(1)_= 5.2645, p < .05).

Exploration time analysis (two-way ANOVA with variance modeling to correct for heteroskedasticity) showed a significant treatment-by-familiarity interaction (χ^2^_(1)_= 4.7689, p < .05); however, animals in both groups explored the novel object significantly more than the familiar one (Fig. S5c; VEH: p < .001; BAY: p < .05). As in the experiment with the immediate injection, these results indicate that NF-kB inhibition produces memory impairment, without completely abolishing memory for the familiar object.

In summary, localized NF-kB inhibition in the dorsal hippocampus impaired persistent recognition memory at the same two critical time points identified for HDAC3 inhibition: immediately and six hours after training.

To further characterize the effect of BAY 11-7082, two groups of cannulated animals underwent a sTR in the NOR task immediately followed by an intra-hippocampal injection of either drug or vehicle. Forty-five min later, brains were fixed, and coronal sections near the site of injection were immunostained for p65 to assess the drug’s effect on NF-kB in the hippocampus (Fig. 4a).

We found a significant treatment-by-region interaction in p65-positive cell density (F_(2,44)_ = 6.7827, p < .01). BAY-treated animals showed significantly fewer p65^+^ cells in the CA1 subregion compared to the control group (Fig. 4e, f; cells/mm^3^ = 49.8 ± 7.61, cells/mm^3^ = 30.2 ± 4.58, p < .01). Surprisingly, the BAY group also showed a significant increase in the number of p65^+^ cells in the CA3 subregion (Fig. S5d, e; cells/mm^3^ = 17.3 ± 5.34, cells/mm^3^ = 34 ± 3.44, p < .05), while there were no changes in the DG (Fig. S5f, g). These results show a reduction in p65-marked cells in the region where BAY 11-7082 was injected, which could account for the deficits observed in memory performance.

## Discussion

In this study, we identified two critical periods during memory consolidation, immediately and 6 hours after learning, during which pharmacological manipulation of HDAC3 or NF-kB affects memory persistence in the NOR task. Systemic inhibition of HDAC3 with RGFP966 enhanced memory persistence, and this effect was accompanied by an increase in nuclear p65 in the CA1 region of the hippocampus. On the other hand, **local inhibition of NF-kB in the dorsal hippocampus impaired memory persistence and was associated with a decrease in the total number of p65-positive cells in CA1. Together, these findings point to a potential functional association between HDAC3 and NF-kB signaling during memory consolidation.**

We first confirmed that general inhibition of class I HDACs -which have predominantly nuclear activity- with NaBut enhances memory in the NOR task in mice, as seen earlier by Stefanko and collaborators [9]. Previous work shows that RGFP136, a class I HDAC inhibitor that preferentially inhibits HDAC3, administered immediately after training, promotes memory persistence in both the NOR and NOL tasks in mice [17]. This evidence points to a role of HDAC3 in the consolidation of persistent forms of memory, and here we showed that systemic inhibition with the specific inhibitor RGFP966 promotes memory retention for at least one week after training in the NOR task. Interestingly, HDAC6 inhibition did not have an effect on memory persistence, although it has been shown to promote long-term memory formation and persistence in the IA task [38]. These discrepancies can be caused by differences in the route of administration, given that the long-lasting effect of TubA was achieved by intrahippocampal, and not systemic, injections. There is also the possibility that HDAC6 fulfills different roles depending on the learning task. Finally, we cannot rule out an effect of systemic HDAC6 inhibition on a 24 h LTM in the NOR task, which remains to be determined.

**We did not observe an effect of systemic HDAC3 inhibition on contextual or cued fear memory under our experimental conditions. This result differs from previous evidence showing that manipulation of HDAC3 activity can influence fear memory [19]. However, the previous study used localized expression of a catalytically inactive HDAC3 mutant, whereas we used systemic pharmacological inhibition, making direct comparisons difficult. Moreover, HDAC3 inhibition may have opposing effects among the different regions involved in fear memory consolidation [58, 59]. Differences in drug penetrance across brain regions [60] and in the optimal dose for this task [18] may also have contributed. Thus, the absence of an effect in our experiment does not necessarily exclude a role for HDAC3 in fear-memory consolidation, but suggests that its contribution may depend on the brain region, memory paradigm, or mode of inhibition. An additional limitation is that fear memory was tested repeatedly at 24 h, 1 week, and 2 weeks after training. Because retrieval can engage reconsolidation processes, previous testing may have influenced subsequent memory performance, making the later time points less suitable as independent measures of the persistence of the original memory.**

**We next evaluated the effect of acute NF-kB inhibition in the hippocampus on NOR memory persistence. As observed with systemic RGFP966 administration, NF-kB inhibition affected memory when administered immediately or 6 h after training, identifying the same temporal windows in which HDAC3 inhibition enhanced memory persistence and closely relating to the two peaks of NF-kB DNA-binding activity found in crabs [36]. A limitation of this experiment is that NF-kB inhibition was assessed using only BAY 11-7082, which may have off-target effects. Thus, an independent NF-kB intervention would strengthen the specificity of these findings.**

**Our findings are also consistent with the broader concept that memory consolidation involves temporally regulated waves of gene expression and protein synthesis after learning [1]. Early experiments in rats showed that there are two phases of protein synthesis in the hippocampus after learning an aversive task: the first occurring around the time of training and the second between 5 and 9 hours later; and that inhibition of either phase impairs LTM formation [30]. Later, these findings were confirmed and expanded by Muller Igaz and collaborators, who demonstrated that inhibition of transcriptional activity also has a detrimental effect on memory consolidation in a biphasic manner [31]. In this context, our findings that RGFP966 administration enhances memory persistence immediately or 6 h after training, but not at 3 or 9 h, while BAY 11-7082 impairs it at those same time points, are consistent with the hypothesis of two phases of critical transcriptional activity.**

Although it has been consistently found that HDAC3 inhibition produces an increase in histone acetylation [17, 18, 20, 61], **previous studies in cell culture have proposed that HDAC3 can also regulate NF-kB activity by deacetylating p65, promoting its interaction with IkB and subsequent nuclear export [25]. Electrophysiological experiments have also provided evidence supporting a relationship between HDAC3 inhibition and NF-kB activation [29]. Based on these findings, we hypothesized that HDAC3 inhibition would increase p65 acetylation and NF-kB nuclear localization during memory consolidation.** Our results did not reveal a clear effect of RGFP966 on p65 acetylation in whole-hippocampus extracts, although they suggest there might be differences in acetylation levels between males and females, for which a larger sample size would be necessary to make the corresponding analysis. It is also important to note that the antibody used to detect acetylated p65 recognizes K310 acetylation only, and other acetyl-lysine residues can also be targeted by HDAC3 [27, 62], so we could have missed part of the effect due to this methodological constraint. In contrast, immunofluorescence analysis revealed an increase in the proportion of nuclear p65 in the CA1 region of the hippocampus after HDAC3 inhibition with RGFP966. **This finding is consistent with our hypothesis that HDAC3 inhibition may promote NF-kB nuclear localization during consolidation and could represent a molecular correlate associated with the memory-enhancing effect of RGFP966**. **However, these immunofluorescence experiments included only three animals per group. Although each animal contributed repeated measurements from multiple brain sections, analyzed with hierarchical mixed-effects models to avoid pseudoreplication, the modest number of biological replicates limits statistical power, and these findings should be interpreted with caution.**

We also observed a reduction in the size of p65 nuclear speckles in the Prh of RGFP966-treated animals, although this difference did not reach statistical significance. Considering there were no differences in intensity in this area, this points to a change in the distribution of the protein, where RGFP966 treatment produces a deaggregation of p65 speckles. This might be related to a previously reported phenomenon where NF-kB (p65 subunit specifically) is sequestered to nucleolar foci under certain conditions as a mechanism to regulate its transcriptional activity [63]. However, further experiments would be needed to determine if this is the case in the context of our experiments. **Moreover, while these findings suggest a relationship between the two pathways, our experiments do not establish a causal link. A combined manipulation of HDAC3 and NF-kB signaling would be required to determine whether NF-kB activity is necessary for the memory-enhancing effects of HDAC3 inhibition.**

We also examined the effect of RGFP966 administration on Zif268 expression and found a significant decrease in protein levels, which was surprising given that *Zif268* is an immediate early gene necessary for object recognition memory consolidation [64] and reconsolidation [65], and that its overexpression enhances spatial recognition [66] and cued fear memory [67]. Although there is evidence that HDAC3 inhibition with RGFP966 promotes Zif268 expression in songbirds [68], this result is from analyzing mRNA levels, which may not be directly correlated with protein levels. One interesting possibility is that Zif268 protein is involved in its own transcriptional repression, as seen previously in cell culture experiments [69]. These findings could explain why protein levels drop while mRNA expression increases. Additionally, we found evidence suggesting sexual dimorphism. Basal sexual differences in Zif268 levels in the medial prefrontal cortex have been previously reported in rats, showing higher levels in males than in females [70]. This supports our findings, in which the vehicle-injected group shows higher Zif268 protein levels in males than in females.

Finally, we examined the effect of hippocampal NF-kB inhibition in p65-stained brain sections and found a decrease in p65-positive cells in CA1, the injection site, which could account for the memory deficits observed. Interestingly, there was an increase in p65-positive cells in CA3, which was unexpected and will require further study.

**Together, our findings provide evidence consistent with a functional relationship between HDAC3 and NF-kB pathways during memory consolidation, while highlighting the need for direct mechanistic experiments to determine whether NF-kB activity mediates the behavioral effects of HDAC3 inhibition on memory persistence.**

## Notes

https://github.com/AgusRobles/Confocal-image-analysis

### Competing Interest Statement

The authors have declared no competing interest.

### Summary of Updates

Attenuated claims of a mechanistic link between HDAC3 and NF-kB throughout; Abstract and Discussion now explicitly state this is not established causally. Title revised to more accurately reflect the results. Corrected terminology, replacing "p65 inhibitor" with "NF-kB pathway inhibitor" (text and Figure legend). Merged NOR and fear conditioning results into one section on RGFP966's behavioral effects; fear conditioning figure moved to supplementary material. Removed BDNF western blot data. Merged BAY 11-7082 behavioral and molecular results into one section; combined corresponding figures; CA3/DG immunofluorescence data moved to supplementary figure. Reorganized Discussion to present behavioral results before molecular evidence; tightened writing throughout. Added rationale for shifting from whole-hippocampus to region/subregion-specific immunofluorescence analysis. Expanded description of the statistical model accounting for nested replicates (animals/sections). Added limitations of small sample size (n=3/group) in immunofluorescence experiments, and potential confound of repeated behavioral testing on memory persistence interpretation in the fear conditioning experiment. Revised immunofluorescence plots to show individual animals with corresponding sections separately from the overall mean. Corrected BAY 11-7082 figure's experimental design: animals received strong training (sTR), not weak training (wTR), before BAY injection.

