## Supplementary information for "HDAC3 inhibitor RGFP966 enhances memory persistence in a biphasic manner and modulates NF-kB nuclear localization"

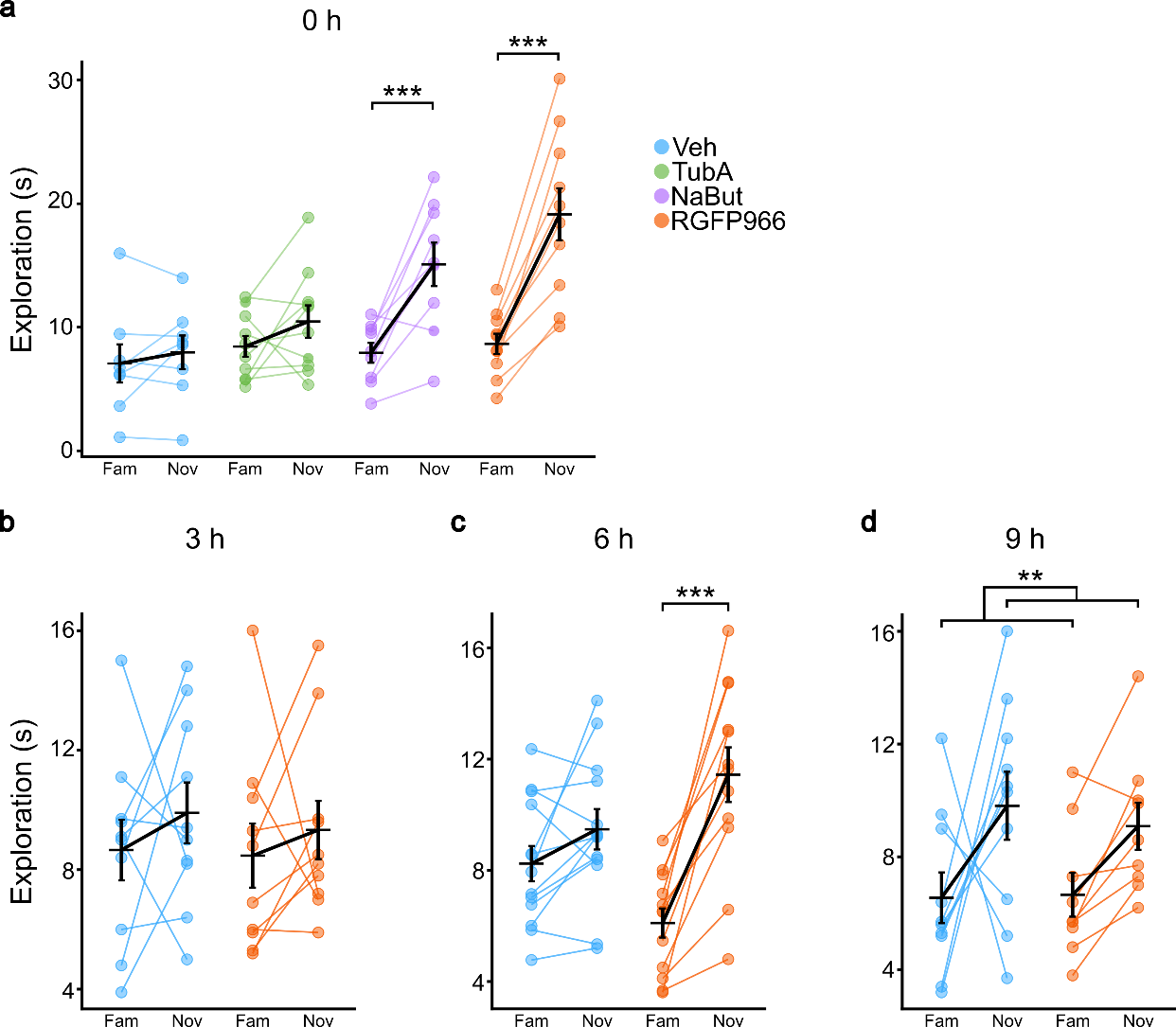

**Fig. S1** Exploration times for familiar and novel objects during testing sessions of animals receiving a wTR in the NOR task followed by i.p. injection of drug or vehicle (**a**) immediately after training; (**b**) 3 hours after training; (**c**) 6 hours after training; and (**d**) 9 hours after training. Mean ± SEM are shown in black. Individual data points are colored by treatment. ** p < .01, *** p < .001; two-way ANOVA, simple pairwise comparisons Tukey-adjusted

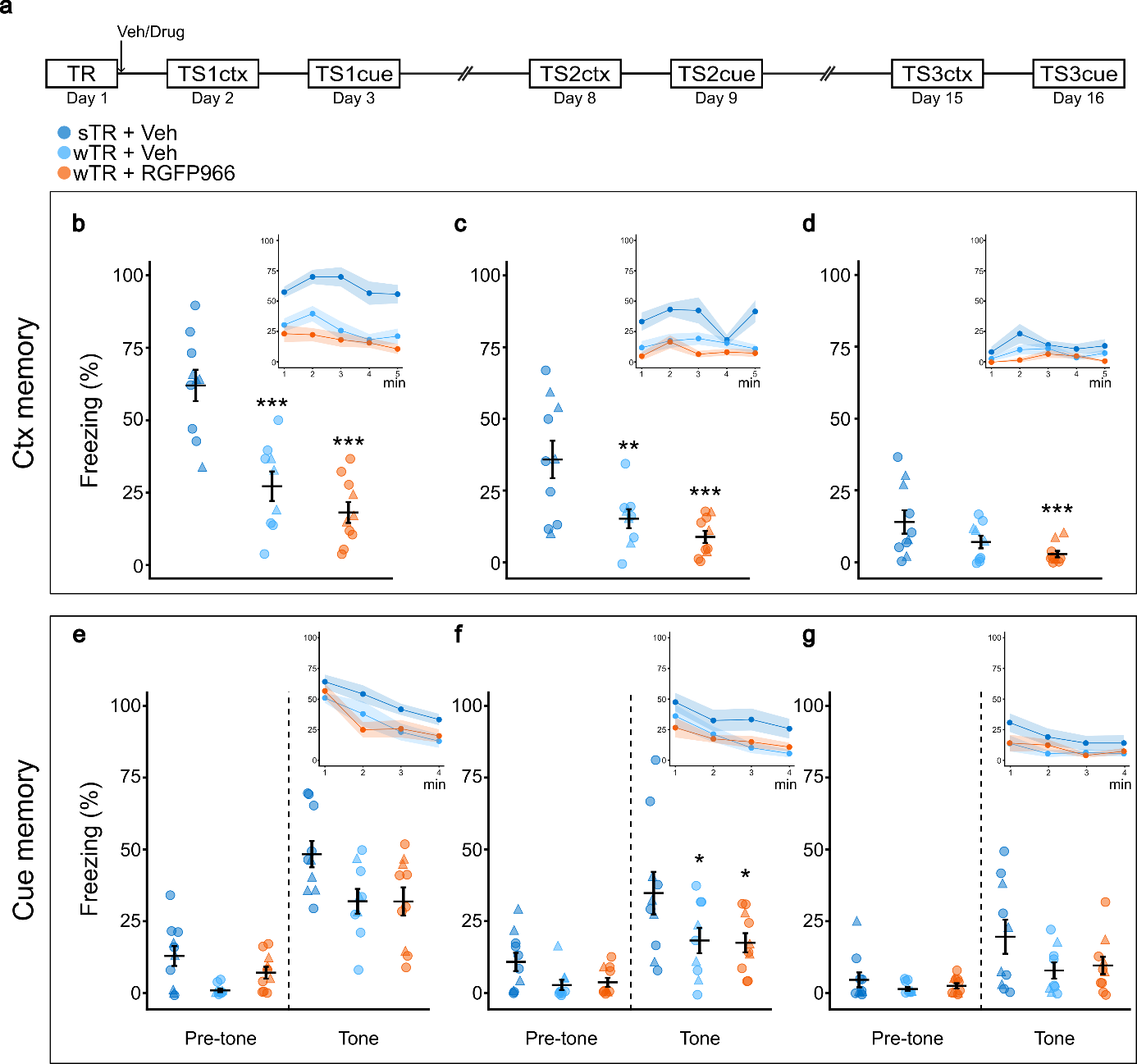

**Fig. S2** Systemic HDAC3 inhibition with RGFP966 does not promote fear memory consolidation of a weak training. (**a**) Experimental design. Animals received either a strong (sTR) or weak training (wTR) immediately followed by an i.p. injection of RGFP966 or vehicle (n_sTR + VEH_ = 10, n_wTR + VEH_ = 9, n_wTR + RGFP966_ = 10). Context (ctx) and tone (cue) memory were tested the next day (TS1), 1 (TS2), and 2 (TS3) weeks after training. (**b-d**) Freezing percentage in the contextual fear memory tests (**b**) 24 h, (**c**) 7 d, and (**d**) 14 d after training. (**e-g**) Freezing percentage in the cued fear memory tests (**e**) 48 h, (**f**) 8 d, and (**g**) 15 d after training. Insets show freezing (%) along each minute of the testing session. Mean ± SEM are shown in black. Individual values are colored by treatment. ▲: females, ⏺: males. Asterisks denote significant differences relative to the sTR group. * p < .05, ** p < .01, *** p < .001. GLMM, type II Wald’s test and Tukey’s multiple comparisons

**Table S1.** p-values of multiple comparisons between the experimental groups in the three testing sessions for contextual and cued memory

|  |  | **Contextual** | **Cued** |
| --- | --- | --- | --- |
| **Session** | **Contrast** | **p-value** | |
| TS1 | sTR VEH - wTR VEH | < .001 | > .05 |
|  | sTR VEH - wTR RGFP966 | < .001 | > .05 |
|  | wTR VEH - wTR RGFP966 | > .05 | > .05 |
| TS2 | sTR VEH - wTR VEH | < .01 | < .05 |
|  | sTR VEH - wTR RGFP966 | < .001 | < .05 |
|  | wTR VEH - wTR RGFP966 | > .05 | > .05 |
| TS3 | sTR VEH - wTR VEH | > .05 | > .05 |
|  | sTR VEH - wTR RGFP966 | < .001 | > .05 |
|  | wTR VEH - wTR RGFP966 | > .05 | > .05 |

**Table S2.** Freezing percentages per group (mean ± SEM) for context (ctx) and cued (cue) testing sessions. TS1: 24/48 h; TS2: 7/8 d; TS3: 14/15 d

| **Session** | **Group** | **Freezing (%) ctx** | **Freezing (%) cue** |
| --- | --- | --- | --- |
| TS1 | sTR  + VEH | 62 ± 5.43 | 48.3 ± 4.58 |
|  | wTR + VEH | 27.2 ± 5.1 | 31.9 ± 4.32 |
|  | wTR + RGFP966 | 18.2 ± 3.6 | 31.9 ± 4.92 |
| TS2 | sTR  + VEH | 35.8 ± 6.53 | 34.8 ± 7.4 |
|  | wTR + VEH | 15.2 ± 3.29 | 18.3 ± 4.44 |
|  | wTR + RGFP966 | 8.83 ± 2.11 | 17.5 ± 3.33 |
| TS3 | sTR  + VEH | 14 ± 4.07 | 19.6 ± 5.92 |
|  | wTR + VEH | 7.04 ± 2.18 | 7.87 ± 2.82 |
|  | wTR + RGFP966 | 2.83 ± 1.11 | 9.58 ± 3.01 |

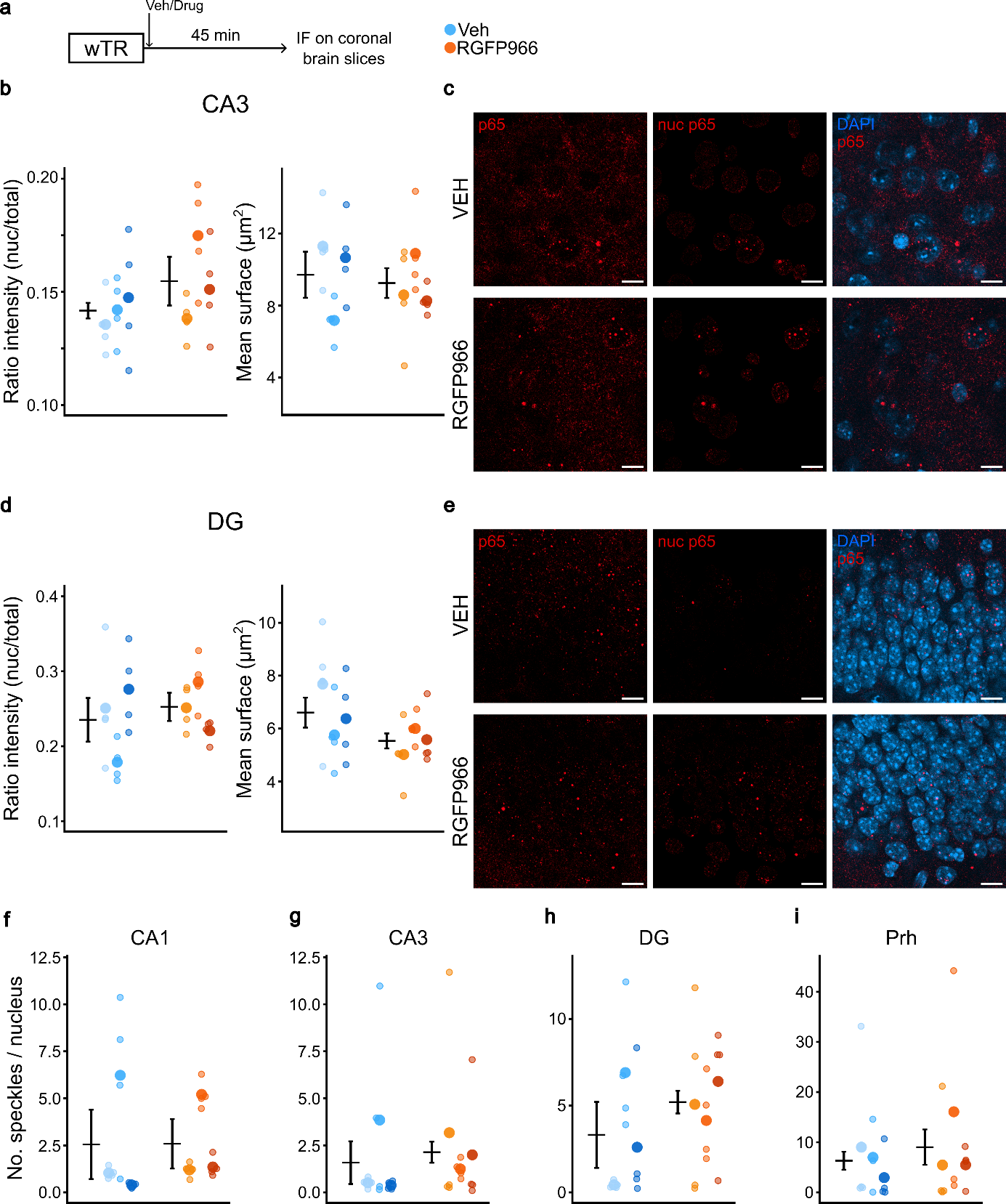

**Fig. S3** Effect of systemic administration of RGFP966 after training on p65 signal in the hippocampus and the Prh. (**a**) Experimental design. Animals received a wTR in the NOR task, followed by an i.p. injection of drug or vehicle. Forty-five minutes later, they were perfused to obtain brain tissue. (**b, d**) (Left) Fluorescence intensity quantification expressed as the proportion of nuclear signal (nuclear p65 / total p65; GLM, type II Wald’s test), and (right) 3D surface area of the p65 speckles inside the nucleus (two-way ANOVA) in (**b**) CA3 and (**d**) DG. (**c, e**) Confocal images of  (**c**) CA3 and (**e**) DG. (**f-i**) Quantification of the number of speckles per nucleus (GLM, type II Wald’s test) in (**f**) CA1, (**g**) CA3, (**h**) DG, and (**i**) Prh. Mean ± SEM are shown in black. Individual data points are colored by treatment. Data from each animal are displayed in separate columns; small dots represent individual brain slices, and the large dot indicates the mean for that animal (n_VEH_ = 3, n_RGFP966_ = 3). Scale bar = 10μm.

**Table S3.** p65 fluorescence ratio, speckle count per nucleus, and speckle surface area across brain regions in VEH- and RGFP966-treated animals. Values are mean ± sem; p-values reflect treatment comparisons within each region

| Region | Group | Ratio | p | # specks | p | Surface (μm^2^) | p |
| --- | --- | --- | --- | --- | --- | --- | --- |
| CA1 | VEH | 0.160 ± 0.008 | < .05 | 4.33 ± 1.86 | > .05 | 8.44 ± 0.88 | > .05 |
|  | RG | 0.188 ± 0.008 |  | 2.59  ± 0.58 |  | 8.85 ± 0.46 |  |
| CA3 | VEH | 0.142 ± 0.005 | > .05 | 1.59 ± 0.90 | > .05 | 9.70 ± 0.75 | > .05 |
|  | RG | 0.155 ± 0.007 |  | 2.15  ± 1.03 |  | 9.24 ± 0.68 |  |
| DG | VEH | 0.235 ± 0.019 | > .05 | 3.32 ± 1.14 | > .05 | 6.61 ± 0.53 | > .05 |
|  | RG | 0.253 ± 0.011 |  | 5.2  ± 1.13 |  | 5.54 ± 0.30 |  |
| Prh | VEH | 0.110 ± 0.009 | > .05 | 6.29  ± 3.06 | > .05 | 10.50 ± 1.41 | = .06 |
|  | RG | 0.114 ± 0.010 |  | 8.40  ± 4.04 |  | 7.15 ± 0.64 |  |

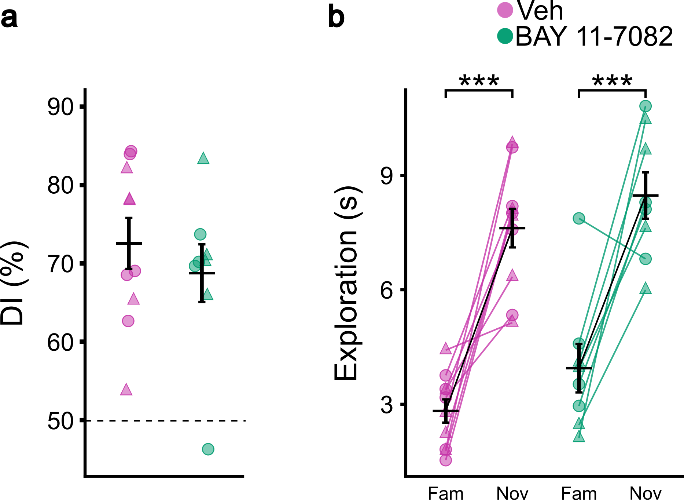

**Fig. S4** Intraperitoneal administration of BAY 11-7082 does not affect memory persistence. (**a**) DI (%) (GLMM followed by type II Wald’s test) and (**b**) exploration times for familiar and novel objects (two-way ANOVA, simple pairwise comparisons Tukey-adjusted) during the testing session of animals receiving an i.p. injection of drug or vehicle immediately after a sTR in the NOR task. Mean ± SEM are shown in black. Individual data points colored by treatment. ▲: females, ⏺: males. *** p < .001

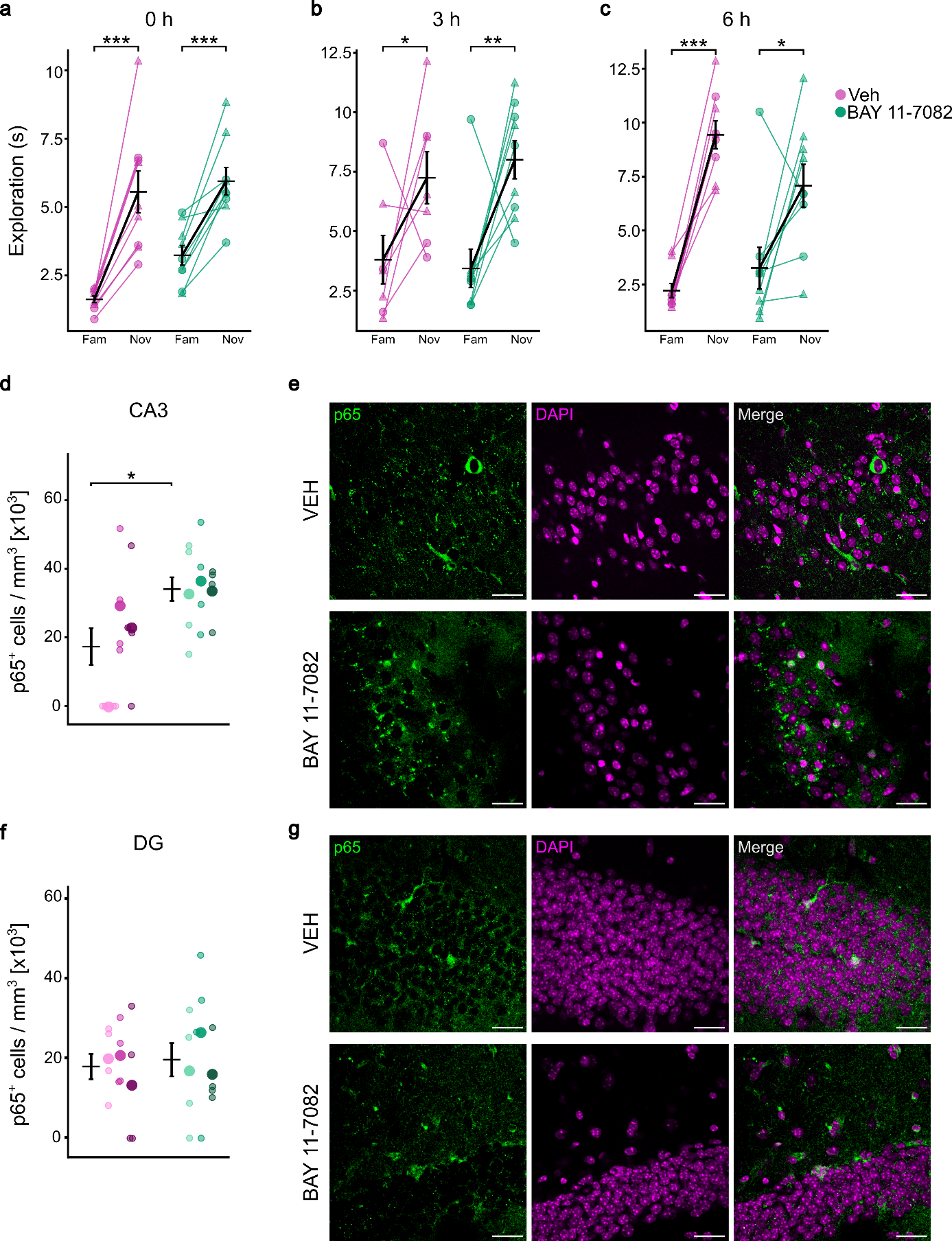

**Fig. S5** Exploration times for familiar and novel objects during testing sessions of animals receiving a sTR in the NOR task followed by intrahippocampal BAY 11-7082 or vehicle injection (**a**) immediately after training, (**b**) 3 hours after training, and (**c**) 6 hours after training (two-way ANOVA, simple pairwise comparisons Tukey-adjusted). (**d, f**) Quantification of the number of positively labeled cells per mm^3^ (two-way ANOVA, simple pairwise comparisons, Tukey-adjusted) in (**d**) CA3 and (**f**) DG. (**e, g**) Confocal images of (**e**) CA3 and (**g**) DG stained against p65 (n_VEH_ = 3, n_BAY_ = 3). Scale bar = 25μm. Mean ± SEM are shown in black. Individual data points are colored by treatment. Data from each animal are displayed in separate columns; small dots represent individual brain slices, and the large dot indicates the mean for that animal. ▲: females, ⏺: males. * p < .05, ** p < .01, *** p < .001
